# Disentangling fluorescence signals from diffusing single molecules by independent component analysis

**DOI:** 10.1101/2025.03.04.641393

**Authors:** Kunihiko Ishii, Miyuki Sakaguchi, Tahei Tahara

## Abstract

Multiparameter measurements offer new possibilities of single-molecule fluorescence experiments for detecting and quantifying heterogeneity in freely diffusing molecules. However, data analysis remains challenging due to limited photon rates from individual molecules. Here, we present independent fluorescence component analysis (IFCA), a universal analytical framework that applies independent component analysis (ICA) to unmix multiparameter fluorescence signals. In this work, we developed an efficient algorithm of ICA for multiparameter photon data based on third-order cumulant tensor decomposition. To validate its performance, we applied IFCA to a static mixture of fluorescent dyes and found that IFCA enables robust separation of signals from five species in a nanomolar mixture solution with a 5 µs binning time. We further analyzed a photon dataset from a FRET-labeled DNA construct using IFCA, demonstrating its capability to quantitatively characterize signals from subpopulations exchanging on a sub-millisecond time scale in a model-independent manner. These findings highlight the high ability of IFCA for quantitative analysis of complex molecular systems exploiting these advantages.

---

Single-molecule fluorescence (SMF) methods, in particular single-molecule Förster resonance energy transfer (smFRET), are invaluable tools in current biophysical/bioanalytical chemistry research for detecting the heterogeneity and fluctuations of molecular structures, which are inaccessible with ensemble measurements.^1, 2^ The major technical challenge in SMF experiments on freely diffusing molecules comes from the intrinsically weak signals that hamper the quantitative analysis of the fluorescence characteristics of constituent species and their relative populations. For deriving maximum information from a limited amount of fluorescence photons, intrinsic properties of fluorescence have been incorporated into SMF experiments, such as excitation/emission wavelengths, emission delay time after pulsed excitation (microtime), and polarization.^3-9^ Photon data obtained with such multiparameter experiments are analyzed with the help of dedicated methods designed for individual experimental configurations, such as photon-by-photon hidden Markov modeling (H^2^MM)^10-12^ and FRET-lines.^13-15^ However, there has been no general approach to photon data analysis that is compatible with an arbitrary combination of fluorescence parameters without assuming a physical model. Moreover, to apply these statistical methods, it is typically necessary to collect at least several tens of photons from each single molecule in a well-isolated condition. This requirement has been limiting the applicability of previous multiparameter SMF methods in terms of sample concentration (∼10 pM)^16^ and time resolution that is determined by temporal bin size or the duration of single-molecule bursts (>∼1 ms).^17^ Also, the brightness of fluorophores needs to be sufficiently high. These factors make it challenging to study molecular events in harsh conditions such as cellular environments.

To solve these problems, we report a new model-free method for unmixing signals in multiparameter SMF experiments based on a multivariate analysis method of signal fluctuations known as independent component analysis (ICA). ICA realizes blind separation of statistically independent components in an observed multiparameter signal by utilizing their non-gaussian property. It has been widely adopted in other fields, e.g., feature extraction, brain imaging, and telecommunications.^18^ Because the fluorescence signals from freely diffusing single molecules recorded with a confocal microscope exhibit Poisson-like non-gaussian behavior,^19^ ICA is naturally expected to be applicable also to SMF data analysis. Here, we implemented an ICA for SMF by using the third-order cumulant tensor^20, 21^ of temporal fluctuations in multiparameter fluorescence signals. We call this analysis IFCA (independent fluorescence component analysis) hereafter.

In this paper, after describing the principle and actual procedures of IFCA, we demonstrate the ability of IFCA by applying it to photon data obtained with multiparameter fluorescence measurements. First, we analyze photon data of a mixture of five fluorescent dyes in nanomolar concentrations. This example illustrates the capability of IFCA in separating individual species and characterizing the fluorescence signal of each species in a high concentration regime. Then, we examine photon data of a FRET-labeled DNA construct that undergoes transitions between two conformational states in a sub-millisecond time scale,^22^ in which transient species in a dynamic system are identified utilizing the microsecond time resolution achievable with IFCA.

## Results and Discussion

### Multiparameter photon data for IFCA

In a multiparameter single-molecule fluorescence measurement (Fig. 1a), a dilute solution of fluorescent molecules is irradiated under a confocal microscope by light pulses of either a single color or multiple colors with temporal interleaving. The emitted fluorescence photons are sorted according to their colors, *λ*_em_, using dichroic mirrors and/or their polarizations using a polarizer and counted by separate detectors. The microtime, *t*, and the excitation wavelength in case of multi-color excitation, *λ*_ex_, of each detected photon are determined from the detection delay time measured with a time-correlated single-photon counting (TCSPC) device. All the photons are also tagged with their absolute arrival times (macrotime), *T*, and stored as multiparameter photon data (Fig. 1a, right panel).^23^ In this data, *λ*_em_ and polarization are indicated by the detector index *c* (for example, *c* = 0 for the donor detector and *c* = 1 for the acceptor detector in a FRET experiment). In case of multi-color excitation, *λ*_ex_ is judged from the value of *t* recorded by the TCSPC module (for example, *t* ≤ *t*_1_ corresponds to a photon emitted after excitation with the first color and *t* > *t*_1_ corresponds to a photon emitted after excitation with the second color). We collectively denote these fluorescence parameters as a one-dimensional variable, *x* = *ct*_0_ + *t*, where *t*_0_ is the number of TCSPC channels, and *x* takes an integer value between 1 and *x*_0_ = *c*_0_*t*_0_ (*c*_0_ is the number of detectors). The goal of IFCA is to obtain the characteristic fluorescence patterns of independent fluorescence components (IFCs) represented as functions of *x*.

**Figure 1.**
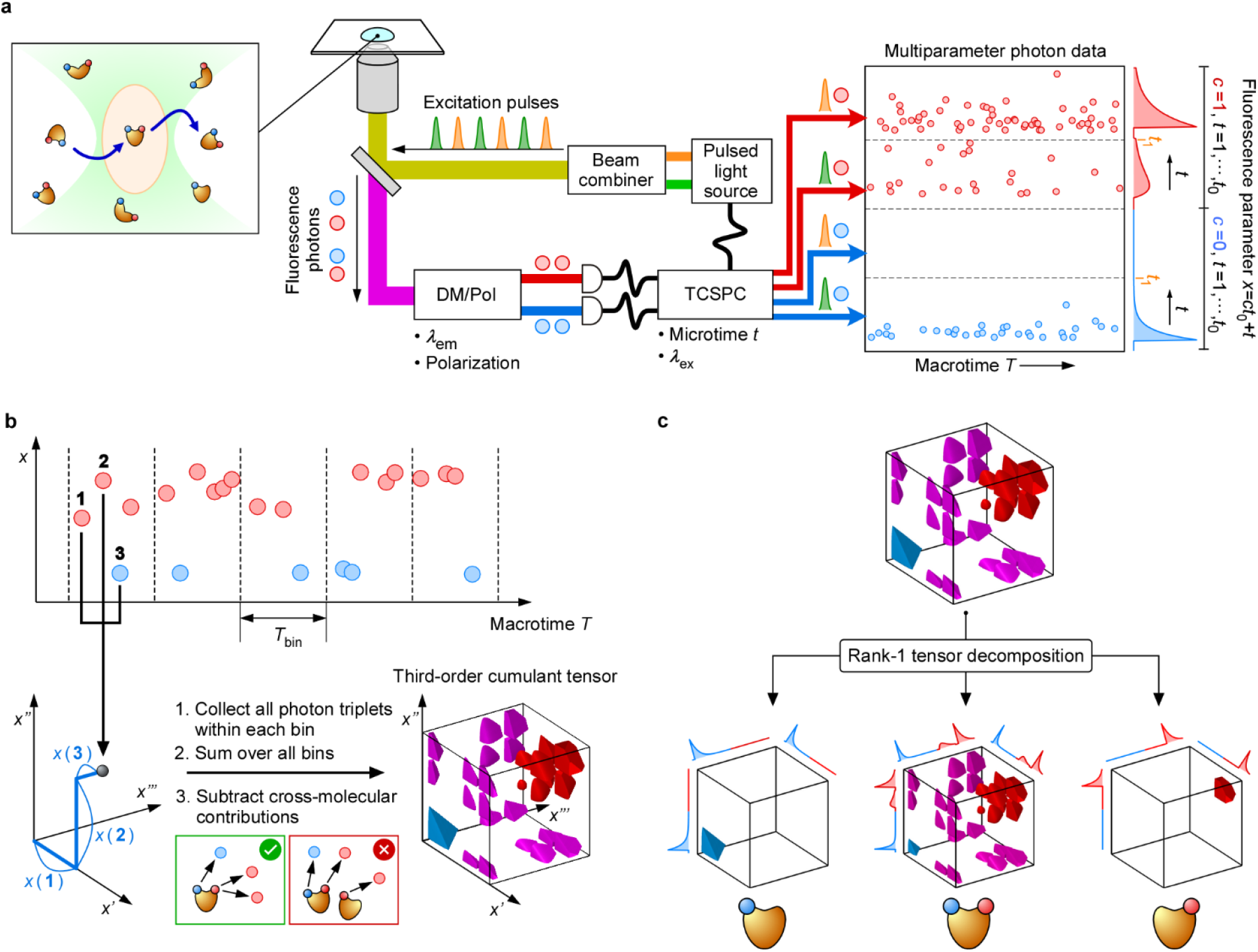
Principle of IFCA. (a) Schematic illustration of a multi-parameter SMF measurement of freely diffusing molecules. DM: dichroic mirror, Pol: polarizer. (b) Construction of a third-order cumulant tensor. (c) Decomposition of a third-order cumulant tensor into a sum of independent fluorescence components.

### Principle of IFCA

IFCA analyzes the fluctuation of fluorescence intensity at channel *x, I*_*x*_(*T*), as a function of macrotime *T*. For this purpose, we utilize third-order cross-cumulants of *I*_*x*_(*T*) that are defined as:^18^

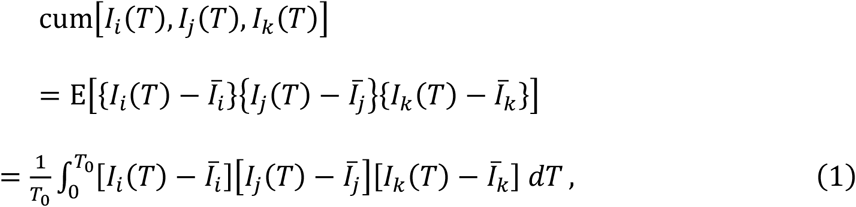

where E[⋅] denotes an expectation value, 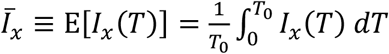 is the time-integrated average of *I*_*x*_(*T*), and *T*_0_ is the total measurement time. The third-order cumulant tensor is defined as a three-dimensional array **𝒯**, whose (*i, j, k*) -element is 𝒯_*ijk*_ ≡ cum[*I*_*i*_(*T*), *I*_*j*_(*T*), *I*_*k*_(*T*)] (Fig. 1b). In IFCA, we exploit an important property of cumulant tensors: the additivity.^18^ That is, a third-order cumulant tensor is decomposed as a sum of multiple rank-1 tensors, each of which is represented as the triple direct product of the vector representing the characteristic fluorescence pattern of a constituent species in the sample (Fig. 1c):

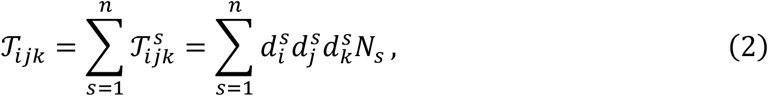

where *n* is the number of species, *N*_*s*_ is the average number of molecules of species *s* in the observation volume, and 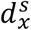 is the characteristic fluorescence pattern of a single molecule of species *s* . For third-order cumulant tensors, there exists another useful property, i.e., uniqueness of decomposition.^24^ Because of this property, the above decomposition of **𝒯** into the contributions of independent sources, **𝒯**^*s*^, is uniquely defined. Based on these properties of cumulant tensors, we realize IFCA in two steps: Construction of the third-order cumulant tensor **𝒯** from multiparameter photon data (Fig. 1b) and decomposition of **𝒯** into independent components (Fig. 1c).

### Procedure of IFCA

#### Construction of the third-order cumulant tensor from multiparameter photon data

To derive third-order cumulants as defined in Eq. 1 from multiparameter photon data, it is necessary to account for the quantum nature of light.^25^ In the fluctuation analyses of fluorescence photons, it is known that the factorial cumulants of photon counts should be employed instead of ordinary cumulants of fluorescence intensity.^19, 26^ In the present case, this can be accomplished by evaluating **𝒯** from photon counts as follows:

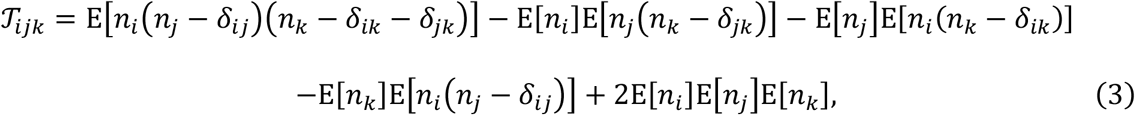

where *n*_*x*_ indicates the photon counts per bin time at channel *x* and *δ*_*xx*′_ is the Kronecker’s delta. Computation of these elements are performed as follows (Fig. 1b).

i. The macrotime axis of the photon data is binned into a preset temporal width, *T*_bin_.
ii. In each bin, photon triplets are searched, and once found, their *x* values are histogrammed in a three-dimensional coordinate space.
iii. The obtained three-dimensional histograms for all bins are summed up and then divided by the total number of bins.
iv. If there are multiple data files recorded in the same measurement condition, the three-dimensional histograms may be averaged over these files.
v. Then, all the possible permutations of the indices (*i, j, k*) are considered, and the obtained six histograms are added up.

It is easy to see that the resultant three-dimensional array corresponds to the first term of Eq. 3. Note that the decrements of *n*_*x*_ by *δ*_*xx*′_ in Eq. 3 are equivalent to avoiding double (or triple) counting of the same photon in a photon triplet in step (ii). The second to fifth terms of Eq. 3 can be evaluated and subtracted from the three-dimensional array by using the first- and second-order cumulant tensors, **v** and **M**:

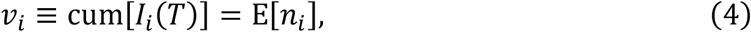

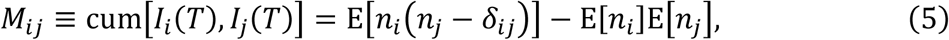

which are computed by histogramming photons and photon pairs, respectively, in a similar manner to the above steps (i)-(v) for deriving 𝒯_*ijk*_. Intuitively, the second to fifth terms of Eq. 3 and the second term of Eq. 5 are corresponding to cross-molecular contributions, i.e., photon triplets (pairs) consisting of photons from different molecules (Fig. 1b), and those involving uncorrelated background counts.^27^ After subtracting these terms in Eq. 3, a third-order cumulant tensor is obtained that can be decomposed into independent components. In practice, it is sometimes necessary for obtaining the best results to consider additional effects due to molecular dynamics within each bin and detector nonideality (Computational Methods 1 in Supporting Information).

#### Decomposition of the third-order cumulant tensor into independent components

Next, we decompose the third-order cumulant tensor into the sum of rank-1 tensors. Though this decomposition is known to be unique provided that all the fluorescence patterns are linearly independent of each other,^24^ the numerical procedure is not as well established as its matrix analogues such as singular value decomposition. Here, we employ a standard method in ICA for preprocessing of higher-order cumulant tensors^28^ utilizing the eigenvalue decomposition (EVD) of the second-order cumulant tensor. In this procedure, we first apply noise normalization across channels to make the noise level uniform over *x*. Because the noise level at a certain channel is proportional to the square root of the counts registered, the normalization factor is set to 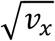 by using the first-order cumulant tensor **v**, i.e., the ensemble-averaged fluorescence pattern. Thus, the raw cumulant tensors **𝒯, M** derived from experimental photon data are scaled as

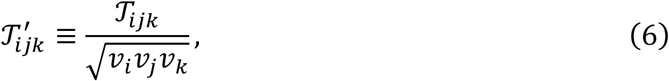

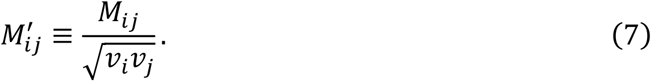

EVD of the scaled second-order cumulant tensor is

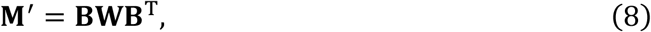

where **B** is a matrix combining the eigenvectors as 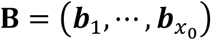 and **W** is a diagonal matrix with the diagonal elements *W*_*p*_ being eigenvalues in the descending order, and superscript T indicates matrix transpose. In the following analysis, only the eigen components with the largest *n* (*n* < *x*_0_) eigenvalues are considered to reduce computational cost and noise. *n* determines the number of detectable IFCs and selected on the basis of the comparison of the first *n* eigenvalues and their associated eigenvectors with the noise components (see Fig. 2c,2d,3c,3d). Using **B, W**, and *n*, it can be shown that an element of the scaled third-order cumulant tensor **𝒯**^′^ is approximated as a sum over *n* independent components (Computational Methods 2 in Supporting Information):

**Figure 2.**
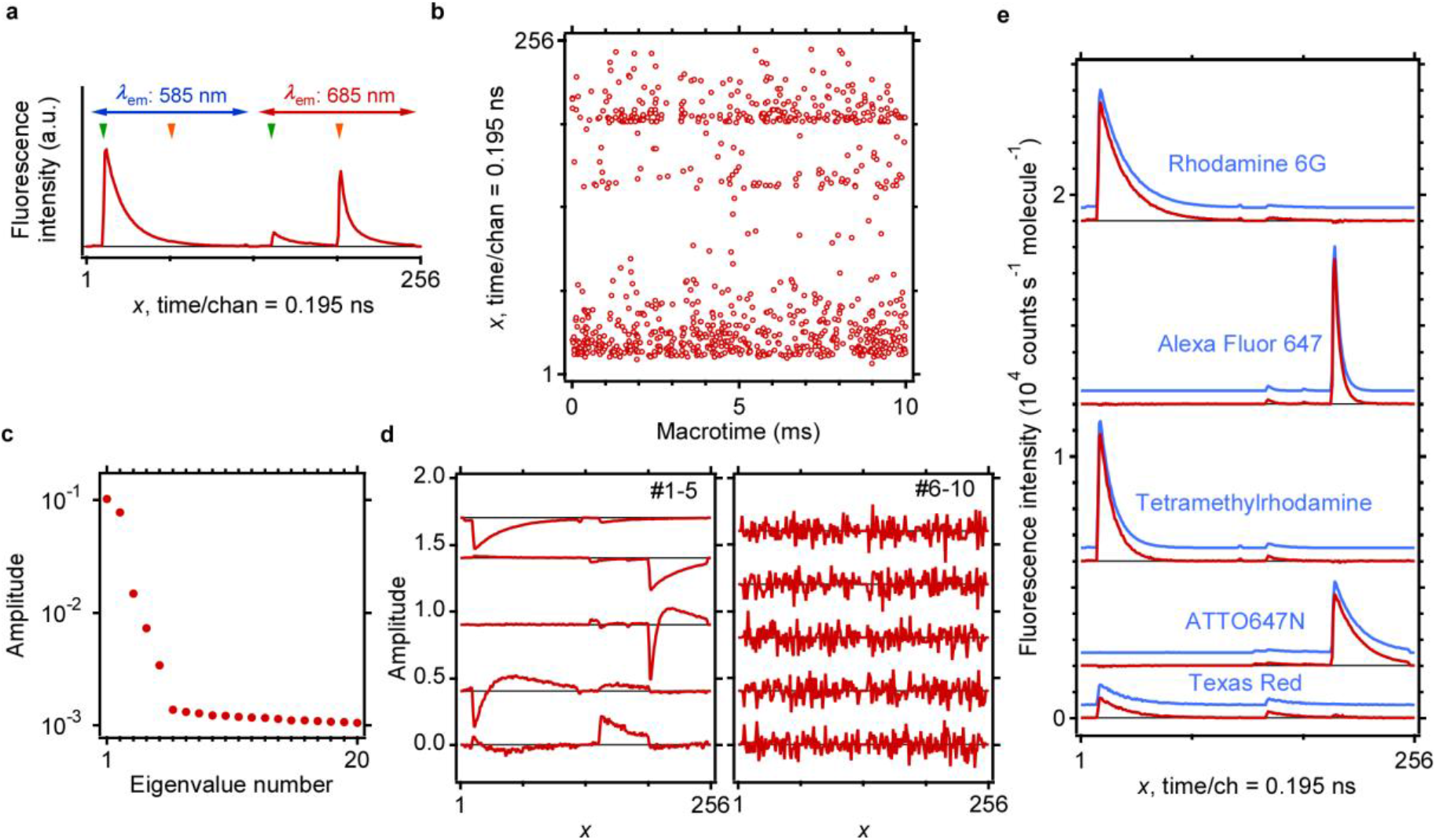
Application of IFCA to a dye mixture. (a) Ensemble-averaged fluorescence pattern of a mixture of five fluorescent dyes. The arrival times of two-color excitation pulses are indicated by downwards arrowheads (green: 525 nm, orange: 640 nm). (b) Representative portion of the photon data of the mixture sample. In this panel, 910 photons are plotted in total. (c) The largest 20 eigenvalues of the scaled second-order cumulant tensor. (d) The eigenvectors associated with the largest 10 eigenvalues (#1-5 (left) and #6-10 (right), from top to bottom). Traces are vertically offset for clarity. (e) Fluorescence patterns of five IFCs retrieved by the IFCA analysis (red). Traces are vertically offset for clarity. Fluorescence patterns of individual fluorophores are shown in blue, which are scaled and vertically offset for comparison.

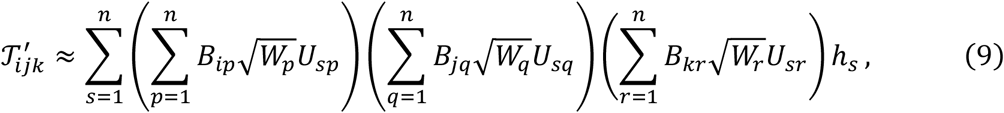

where *h*_*s*_ is a coefficient representing the contribution of the *s*-th independent component, which contains information of the average number of molecules in the observation volume, and *U*_*sp*_, *U*_*sq*_, *U*_*sr*_ are elements of an *n* × *n* orthogonal matrix **U**. Hence, the goal of IFCA is to determine *h*_1_, ⋯, *h*_*n*_ and **U** by comparing Eq. 9 and the experimentally obtained cumulant tensor. To carry out this analysis efficiently, the dimension of the third-order cumulant tensor is reduced by multiplying the eigenvectors:

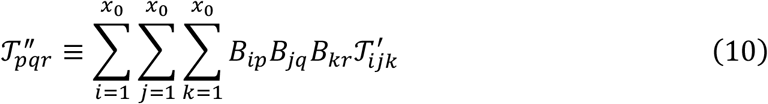

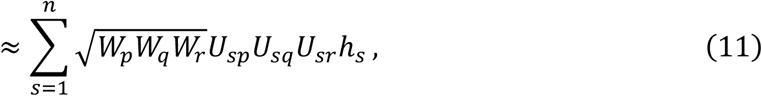

where 1 ≤ *p, q, r* ≤ *n*. With this conversion, the original dimension of 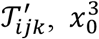 (e.g., *x*_0_ = 256), is reduced to *n*^3^, where *n* is typically smaller than 10. *h*_1_, ⋯, *h*_*n*_ and **U** are determined by fitting the experimentally obtained **𝒯**^″^ (Eq. 10) with Eq. 11 by using *h*_1_, ⋯, *h*_*n*_ and **U** as fitting parameters (Figs. S2, S3 in Supporting Information). Fitting involving an orthogonal matrix as a fitting parameter can be performed by a Jacobi-type algorithm^28^ (Computational Methods 3 in Supporting Information). Finally, the fluorescence patterns of IFCs are reconstructed from the obtained parameters (Computational Methods 2 in Supporting Information):

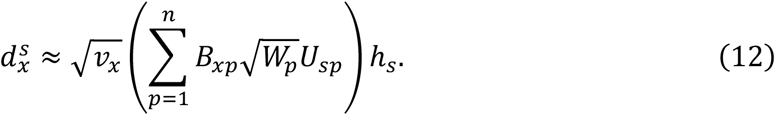

Here, the factor 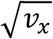 is multiplied to compensate for the noise normalization (Eqs. 6 and 7). This 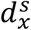 represents fluorescence pattern of a single molecule of component *s*, and its sum over *x* corresponds to molecular brightness. Note that the shape of 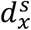 potentially depends on the choice of *n*.

The concentrations of fluorescent species representing IFCs are evaluated as the average numbers of molecules in the observation volume, *N*_*s*_, which is obtained as (Computational Methods 2 in Supporting Information)

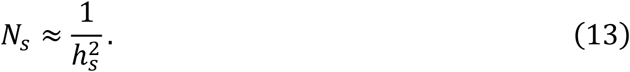

Note that this is the average number of molecules in the observation volume having a well-defined boundary. In SMF measurements under confocal microscope such as fluorescence correlation spectroscopy (FCS), the observation volume is often approximated by a 3-D gaussian profile. In that case, the value obtained with Eq. 13 differs from that defined in FCS, 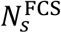, as:^19^

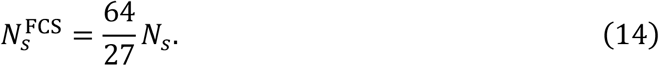

Details of the IFCA analysis on actual photon data are described in Supporting Information, Computational Methods 4.

### Application to a mixture of fluorescent dyes

As the first example, we applied IFCA to experimental photon data of a static mixture of five fluorescent dyes. A mixture solution of tetramethyl rhodamine (TMR), rhodamine 6G (R6G), Alexa Fluor 647 (Alexa647), ATTO647N, and Texas Red (TR) was prepared with a concentration of 1-10 nM for each dye (Experimental Section 1 in Supporting Information). These dyes have different absorption/emission wavelengths and/or fluorescence lifetimes.

Multiparameter fluorescence measurement was performed on this mixture solution with a confocal microscope equipped with pulsed-interleaved excitation (PIE)^7, 29^ and TCSPC (Experimental Section 2 in Supporting Information).^30^ Figure 2a shows the fluorescence pattern of the mixture solution integrated over the entire acquisition period (3313 sec.). This pattern is essentially the sum of contributions from individual dyes having specific fluorescence patterns. The multiparameter photon data obtained in this experiment (Fig. 2b) exhibits two notable features. First, photons seem to be detected mostly at random and distributed uniformly. In other words, no time regions are clearly discernible that exhibit locally dense photons. The lack of such ‘burst-like’ features is due to the relatively high dye concentrations (nM), making the observation volume almost always occupied with at least one molecule. Such a high concentration regime is usually not compatible with ordinary single-molecule experiments, where separation of individual molecules is of paramount importance. Secondly, the average interval of photon arrival times is about 10 µs. If we performed a typical smFRET experiment with such a count rate, it would take ∼1 ms to collect sufficient number of photons to derive a reliable FRET efficiency. Therefore, even if a lower concentration were employed in this experiment, it would be impossible to probe transient states with a microsecond lifetime.

Figures 2c,d show the results of EVD analysis (Eq. 8) of the scaled second-order cumulant tensor made from the recorded photon data employing the temporal binning width of 5 µs. In the eigenvalue plot (Fig. 2c), it is clearly seen that the first five largest eigenvalues are beyond the noise level. Correspondingly, the associated eigenvectors of these eigenvalues (Fig. 2d, #1-5) display meaningful features, while other eigenvectors (#6-10) are dominated by noise. Hence, we selected the first five eigenvectors for reducing the third-order cumulant tensor (Eq. 10), and then the reduced tensor was fitted using Eq. 11 to obtain *h*_1_, ⋯, *h*_5_ and **U**. The fitting results satisfactorily match with the elements of the reduced tensor (Fig. S2 in Supporting Information), justifying the use of fitting algorithm employed in this work.

The five IFCs recovered by Eq. 12 are compared with reference patterns determined by individually measuring solutions containing only one of the five dyes (Table 1). In this table, the cosine similarity between a recovered IFC, *s*, and a reference pattern, *r*, are evaluated as

**Table 1.**
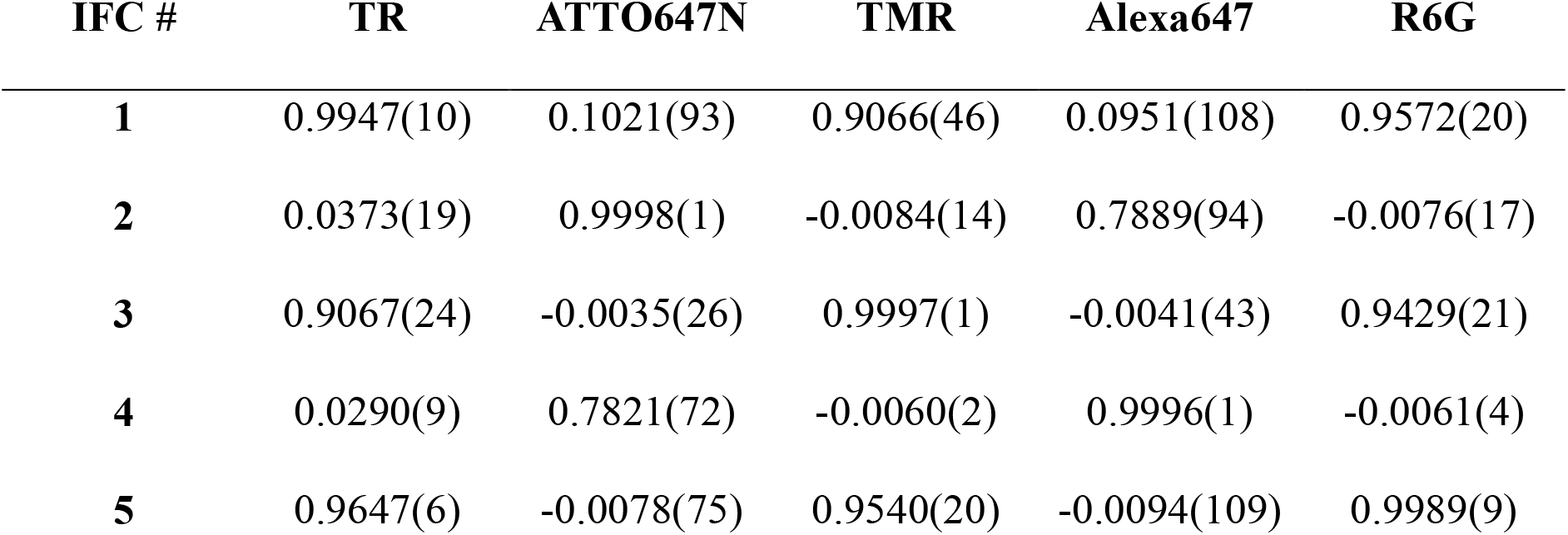
Cosine similarities between the IFCs retrieved by IFCA and reference fluorescence patterns of the five fluorophores. Values in parentheses are the standard deviations (multiplied by 10^4^) estimated from three measurements.

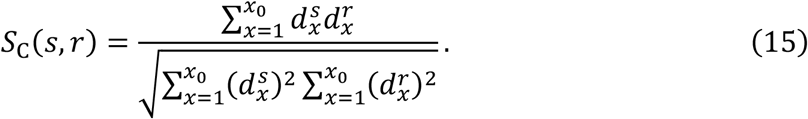

For each of the recovered IFCs, a reference pattern exhibiting a high similarity (*S*_c_ > 0.99) is found. According to Table 1, we made assignments of IFCs #1, #2, #3, #4, and #5 as TR, ATTO647N, TMR, Alexa647, and R6G, respectively. The fluorescence patterns of these IFCs and assigned reference patterns (Fig. 2e) are visually almost indistinguishable, confirming the successful decomposition of IFCs using IFCA. The average numbers of molecules in the observation volume derived by Eq. 13 are *N* = 0.343 (TR), 0.095 (ATTO647N), 0.089 (TMR), 0.088 (Alexa647), and 0.070 (R6G). The relative concentrations estimated from these values are consistent with the actual concentration (5 nM for TR and 1 nM for ATTO647N, TMR, Alexa647, R6G) within experimental error in sample preparation. Furthermore, the fluorescence patterns of IFCs and their relative concentrations are reproducible for three measurements (Fig. S1f,g in Supporting Information).

We have also examined the dependence of IFCs on the number of components (*n*) selected for reducing the third-order cumulant tensor (Eq. 10). The results for setting *n* = 4 to 6 are presented in Fig. S1a-c in Supporting Information. When *n* is decremented (*n* = 4), the IFC assignable to TR disappears, while when *n* is incremented (*n* = 6), an unphysical noisy component appears. These are naturally expected behaviors and regarded as representing the practicality of IFCA. On the other hand, the obtained fluorescence patterns are essentially invariable on extending the binning width *T*_bin_ up to 50 µs (Fig. S1d, e in Supporting Information), which implies that the result for *T*_bin_ = 5 µs is reliable, as this measurement was performed on a static mixture sample, and hence, there should be no binning-width dependence. At the same time, an increase in the *N* values and concomitant decrease of the amplitudes of fluorescence intensity, which corresponds to the molecular brightness, are observed on increasing *T*_bin_. This behavior is ascribed to the effect of molecular diffusion across the observation volume.^31^ Notably, the binning width employed in this analysis, *T*_bin_ = 5 µs, is even shorter than the average photon interval, ∼10 µs (Fig. 2b), which is in a stark contrast with conventional methods of smFRET analysis.

### Application to a FRET-labeled DNA construct

In the second example, we focused on application of IFCA to a molecular system undergoing dynamic transitions between conformational states distinguished by their FRET efficiencies. For this purpose, we analyzed photon data sets that were obtained by Schröder et al.^22^ and deposited on a public repository.^32^ The data contain photons recorded with two-color TCSPC-FCS measurements on a single-stranded DNA, which was labeled with Cy3B as a FRET donor and tethered on a small DNA origami structure as illustrated in Fig. 3a. In addition to this donor-labeled single-stranded DNA (*pointer* strand), the DNA origami structure was attached with two short single-stranded DNA (*staple* strands) having identical 5-nt sequences complementary to the end of the pointer strand. With this design, the pointer strand may form a duplex with either of the two staple strands with almost equal probability. The FRET acceptor (ATTO647N) was fixed on the DNA origami structure near one of these two staple strands, so that switching between the two states could be detected in an equilibrium condition through the change of FRET efficiency. The relaxation time constant between these two states, high-FRET (HF) and low-FRET (LF), was reported to be *τ* = (1/*τ*_HF→LF_ + 1/*τ*_LF→HF_)^−1^ = 220 ± 20 µs.^22^ The entire DNA construct was freely diffusing in a buffer with a concentration of 100 pM.

**Figure 3.**
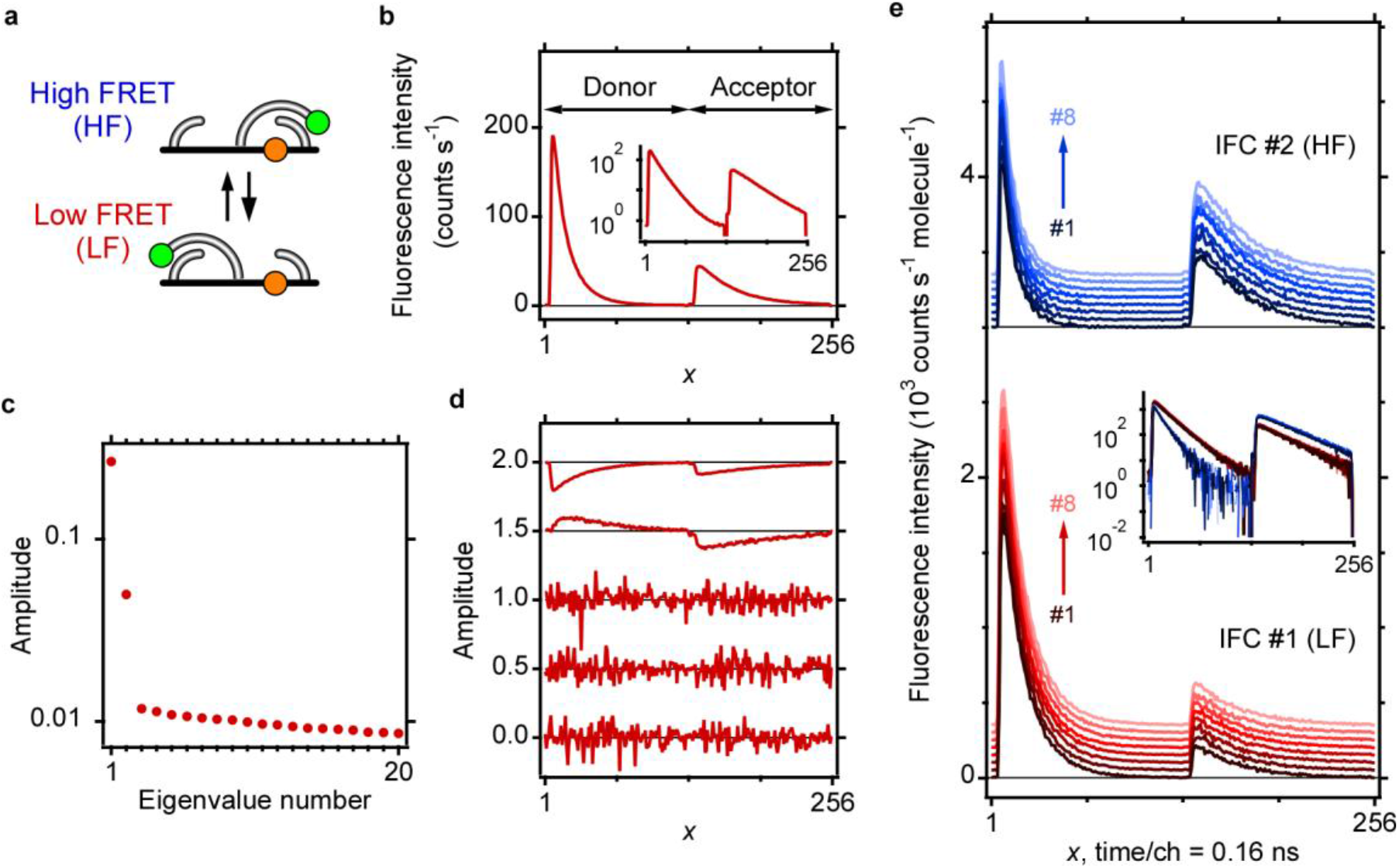
Applications of IFCA to a FRET-labeled DNA construct. (a) Schematic illustration of the FRET-labeled DNA construct. Green and orange circles: donor and acceptor dyes, black bars: DNA origami structure. (b) Ensemble-averaged fluorescence pattern. Inset: The same pattern in log scale. (c) The largest 20 eigenvalues of the scaled second-order cumulant tensor. (d) Eigenvectors associated with the largest five eigenvalues (from top to bottom). Traces are vertically offset for clarity. (e) Fluorescence patterns of IFCs retrieved by the IFCA analysis. Two IFCs obtained for the eight data sets are plotted separately with vertical offsets for clarity. Inset: All IFCs plotted together in log scale.

Figure 3b shows a concatenated fluorescence pattern of the donor fluorescence (*x* = 1-128) and acceptor fluorescence (*x* = 129-256) integrated over 600 s. The decay time of the donor fluorescence and the ratio of integrated intensities of the donor (*I*_D_) and acceptor (*I*_A_) fluorescence reflect the FRET efficiency of the labeled DNA. This ensemble pattern is the superposition of the HF and LF patterns, which is characterized by faster donor decay with higher acceptor-to-donor ratio, and slower donor decay with lower acceptor-to-donor ratio, respectively. To separate these contributions, we applied IFCA to the raw photon data by employing a binning width *T*_bin_ = 20 µs that is much shorter than the relaxation time scale.

Being consistent with the two-state model in Fig. 3a, two significant eigen components were identified in an EVD analysis (Figs. 3c,d). It indicates that a coexisting impurities such as donor-only population is negligible, in agreement with the result of a strict single molecule analysis by Schröder et al.^22^ Therefore, we set *n* = 2 and performed the dimensionality reduction and fitting of the third-order cumulant tensor using Eqs. 10 and 11 (Fig. S3 in Supporting Information). The obtained fluorescence patterns (Fig. 3e) are undoubtedly assignable to the LF (IFC #1) and HF (IFC #2) states due to rapid donor decay and relatively high acceptor intensity of IFC #2. This result is highly reproducible over eight data sets, allowing us to precisely determine the mean donor fluorescence lifetimes (τ_donor_) and the apparent FRET efficiencies (*E* = *I*_A_/(*I*_D_ + *I*_A_)). These values were determined as 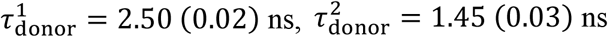 and *E*^1^ = 0.179 (0.005), *E*^2^ = 0.594 (0.011) for the two IFCs, where values in parentheses are standard deviations. The concentrations were obtained for these IFCs as *N*_1_ = 0.0238 (0.0016) and *N*_2_ = 0.0205 (0.0015), leading to the equilibrium constant *K* = [HF]/[LF] = *N*_2_/*N*_1_ = 0.87 (0.08). Deviation of *K* from unity is consistent with the previous work by Schröder et al.,^22^ who suggested that LF is slightly more stable in similar DNA constructs.

In the present analysis, the binning width of 20 µs allows us to distinguish two forms interconverting on the sub-millisecond time scale. Assuming an exponential dynamics with the relaxation time constant of *τ* = 220 µs, the effect of interconversion in the retrieved IFCs with *T*_bin_ = 20 µs is estimated to be ∼4%. Thus, this analysis highlights the feasibility of IFCA interrogating dynamic FRET systems with a microsecond time resolution. The results are generally consistent with the previous work by Schröder et al.^22^ Moreover, IFCA provides additional sources of information such as number of IFCs, acceptor intensity, and the precise shapes of decay curves. Interestingly, the donor decays are not exactly single exponential but double exponential functions are required to reproduce them (Fig. S4 in Supporting Information). Such a detailed analysis is made feasible by the model-free nature of IFCA.

## Discussion

The two applications explored in this work represent the high potential of IFCA in resolving mixed fluorescence signals: The first example of the dye mixture illustrates accurate and robust separation of five IFCs and determination of their characteristic fluorescence patterns and relative concentrations on a nanomolar range. The second application to the FRET-labeled DNA construct demonstrates the high time resolution of IFCA to precisely identify microsecond transients and its model-free nature, enabling detailed analysis of the obtained fluorescence patterns. The high time resolution of IFCA and its compatibility with relatively high sample concentrations originate from its basis on a correlation analysis. It is analogous to FCS, in which a nanosecond time resolution can be achieved for observing molecular dynamics.^33^ IFCA collects photon triplets emitted from individual molecules and analyzes their statistical properties (Fig. 1b). Because only three photons are required per molecule—far fewer than ∼100 photons typically needed for smFRET—short binning times down to a few microseconds can be employed. On the other hand, the advantage at high-concentrations stems from the ability of IFCA to accurately eliminate cross-molecular contributions caused by multiple molecules coexisting within the observation volume. This is accomplished through the construction of cumulant tensors (Eqs. 3 and 5), which automatically subtract uncorrelated backgrounds including cross-molecular effects.^27^ Such automatic subtraction is also convenient for removing background signals originating from water Raman scattering, autofluorescence of optical elements, and detector dark counts, without resorting to additional control measurements (compare insets of Fig. 3b and 3e).

Existing methods for multiparameter SMF analysis of freely diffusing molecules can be categorized into strict single-molecule methods and correlation-based approaches. The former, such as H^2^MM^10-12^ and FRET lines,^13, 14^ require low concentrations (∼10 pM) and high molecular brightness (∼100 photons per bin/burst). Moreover, when applied to study microsecond transients, it is necessary to assume appropriate physical models, such as the relation between the donor/acceptor ratio and fluorescence lifetime^15^ or Markovian property of state transitions. In contrast, IFCA is entirely model-independent except for the assumption of statistical independence of individual components and is compatible with nanomolar concentrations and low brightness (three photons per bin). A representative correlation-based method is two-dimensional fluorescence lifetime correlation spectroscopy (2D FLCS).^34, 35^ Since 2D FLCS is based on FCS, it shares the advantages of IFCA in sample concentration (nM) and time resolution (microseconds).^36^ However, 2D FLCS solely relies on differences in fluorescence lifetimes for component separation. Hence, it is not straightforward to fully utilize information obtainable with multiparameter measurements, though a few techniques have been developed that incorporate additional detection channels.^30, 37^ Therefore, in terms of component separation, IFCA can be regarded as a generalization of 2D FLCS to model-free analysis via employing third-order cumulant tensors. Other correlation-based approaches utilizing higher-order cumulants have been reported by Müller et al.;^19, 31, 38^ however, they were restricted to photon data of up to two detection channels. In contrast, the diagonalization of third-order cumulant tensors introduced in this work enables the separation of general multiparameter signals.

Although IFCA is highly effective for separating multiple components, there are a few specific situations warranting caution. First, fluorescence patterns of two or more components are not always linearly independent. In such cases, the number of mutually orthogonal eigenvectors obtained with EVD of the scaled second-order cumulant tensor (Eq. 8) does not correspond to the true number of components, thereby precluding the application of ICA. Examples include oligomer formation accompanied by changes in brightness without altering fluorescence patterns, and FRET measurements with coexisting donor-only, acceptor-only, and zero-FRET species.^30^ Dedicated analytical protocols are required to be established for these cases. Second, when rapid interconversion occurs between components, it would be difficult to employ a sufficiently short binning width (e.g., less than 10% of the time scale of the dynamics) to minimize the dynamical effect. If the binning width approaches the time scale of dynamics, IFCA yields mixed fluorescence patterns rather than distinct components. Consequently, investigations of fast dynamics require methods to estimate mixing effects accurately, e.g., through systematic analysis of binning-width dependence. Finally, when the number of components *n* is large, it would become challenging to achieve a sufficiently low noise level to extract *n* significant components via EVD (Eq. 8). In such cases, alternative approaches that do not rely on dimensionality reduction are necessary.

The advantages of IFCA demonstrated in this work are expected to enable a broad range of novel measurement modalities. Its high time resolution can be exploited not only for characterizing short-lived transient species but also for measurements under continuous flow in microfluidic systems, where the observation window is constrained by the brief passage time through the detection volume.^39, 40^ Compatibility with high sample concentrations facilitates efficient data acquisition and allows detection of complex formation with moderate (∼nM) affinity via intermolecular FRET.^41, 42^ Most importantly, IFCA is entirely independent of any specific theoretical model for reproducing fluorescence patterns. As a result, it can be applied to arbitrary combinations of fluorescence parameters, i.e., excitation/emission wavelengths, emission delay time, and polarization. The model-free nature enables direct integration with advanced multiparameter SMF techniques such as polarization-resolved measurement^8^ and multi-color FRET.^43^ Beyond single-molecule studies, the same framework is potentially applicable to multiparameter single-particle^44^ or single-cell spectroscopic measurements such as spectral flow cytometry.^45^

## Conclusion

In this work, we applied ICA to multiparameter SMF photon data from freely diffusing molecules and introduced IFCA as a novel analytical framework, based on the decomposition of third-order cumulant tensors generated from photon data. To assess its performance, we analyzed two representative experimental datasets: a static dye mixture and a dynamic FRET-labeled DNA construct. The results demonstrate that IFCA enables robust separation of fluorescence signals from multiple species in samples with concentrations up to the nanomolar range, achieving time resolutions on the order of several microseconds, all within a fully model-free approach. These findings establish IFCA as a versatile and powerful method that holds significant promise for advancing the scope and precision of SMF experiments, paving the way for the quantitative analysis of complex molecular systems.

## Supporting Information

The authors have cited additional references within the Supporting Information.^46-48^

## Data availability

Data for the dye mixture experiment have been deposited at Zenodo and are publicly available at https://doi.org/10.5281/zenodo.14869092. Data for the FRET-labeled single-stranded DNA construct are made publicly available by Schröder et al. and accessible at https://doi.org/10.5281/zenodo.7254095.

## Code availability

A Python module for IFCA analysis has been deposited at Zenodo and is publicly available at https://doi.org/10.5281/zenodo.14869092.

## Acknowledgments

This work was supported by JSPS KAKENHI Grant Number JP23K21102 (to K. I.).

## Notes

https://doi.org/10.5281/zenodo.14869092

### Competing Interest Statement

The authors have declared no competing interest.

### Summary of Updates

The contents are updated by adding detailed descriptions of the results.

