## Supplementary Methods and Figures for "Disentangling fluorescence signals from diffusing single molecules by independent component analysis"

### Table of Contents


**Experimental Section**

1. Sample preparation of a mixture of fluorescent dyes
2. Multiparameter fluorescence measurement

**Computational Methods**

1. Effects of molecular dynamics and detector nonideality
2. Derivation of fluorescence patterns and concentrations
3. Jacobi-type algorithm for determination of $h_1, \cdots, h_n$ and $\mathbf{U}$
4. Details of IFCA analysis

**Supporting Data**

Figure S1. Validation of the stability of IFCA analysis on the mixture sample

Figure S2. Fitting analysis of the reduced third-order cumulant tensor: Dye mixture

Figure S3. Fitting analysis of the reduced third-order cumulant tensor: FRET-labeled DNA construct

Figure S4. Fitting analysis of the donor decays in two IFCs from the FRET-labeled DNA construct

#### Experimental Section

##### 1. Sample preparation of a mixture of fluorescent dyes

We measured a mixed dye solution of 5(6)-carboxy tetramethyl rhodamine (TMR, BioReagent), rhodamine 6G (R6G, Sigma-Aldrich), Alexa Fluor 647 carboxylic acid tris(triethylammonium) salt (Alexa647, Invitrogen), ATTO647N free COOH (ATTO-TEC), and Texas Red-X succinimidyl ester mixed isomer (TR, Invitrogen). The mixture sample was prepared in concentrations of 1 nM, 1 nM, 1 nM, 1 nM, 5 nM for TMR, R6G, Alexa647, ATTO647N, TR, respectively, in 50 mM PBS buffer (pH7.2) containing 0.01% Triton X-100. Reference samples containing only single dyes were prepared in the same concentration and buffer conditions as the mixture sample.

##### 2. Multiparameter fluorescence measurement

We performed an experiment using a home-built multiparameter fluorescence measurement setup. The apparatus is based on the PIE scheme[1, 2] and essentially the same as that was reported previously.[3, 4] The excitation light source is a supercontinuum laser (Fianium SC-400-4, 40 MHz). The visible part of its output was split into two by a dichroic mirror (Thorlabs DMLP567) and used as short- and long-wavelength excitation pulses. They passed through bandpass filters (Semrock, FF01-525/30-25 & FF01-640/14-25) and the timing were adjusted by adding an optical delay to the long-wavelength excitation pulses using a 1-m long single-mode fiber (Thorlabs P1-460B-FC-1), before recombined by another dichroic mirror (Thorlabs DMLP567) and a single-mode fiber (Thorlabs P5-460B-PCAPC-1). They were introduced into a microscope (Nikon Eclipse Ti) and focused onto a sample solution through a water-immersion objective lens (Nikon CFI Plan Apo IR 60×, N.A.=1.27). The solution was filled in a sample chamber made of two coverslips and a silicone spacer. Fluorescence light emitted from the sample was collected with the objective and separated from the scattering of excitation light by a dual-band dichroic mirror (Chroma ZT532/640rpc). After passing through a multi-mode fiber (Thorlabs FG050LGA, core diameter = 50 μm), which acted as a confocal pinhole, short- and long-wavelength fluorescence were separated by a dichroic mirror (Chroma Technology ZT633rdc) and detected by two photon-counting detectors (Becker & Hickl HPM-100-40-C) after passing through bandpass filters (Semrock FF01-585/40-25 & FF02-685/40-25). For each photon detection event, the detector ID was tagged by a routing module (Becker & Hickl HRT-41) and the excitation-emission delay time (microtime) was measured by a TCSPC module (Becker & Hickl SPC-130-EM) in the FIFO mode, i.e., the absolute arrival time (macrotime) was also recorded. The list of macrotime, microtime, and detector ID was saved in the native file format of Becker & Hickl (.spc).

#### Computational Methods

##### 1. Effects of molecular dynamics and detector nonideality

Eqs. (3-5) in the main text assume that the fluorescence intensity $I_x(T)$ does not fluctuate within each bin. Therefore, in principle, $T_{\mathrm{bin}}$ should be short enough as compared with molecular dynamics such as intramolecular conformational transitions or diffusion across the observation volume. However, such a short $T_{\mathrm{bin}}$ may deteriorate the performance of IFCA because it reduces the number of usable photon triplets. Practically, it is recommended to use a $T_{\mathrm{bin}}$ value either sufficiently shorter than or much longer than the time scale of intramolecular dynamics, so that the dynamical effect is well resolved or entirely averaged out, respectively. In this sense, $T_{\mathrm{bin}}$ determines the time resolution to detect transient species. On the other hand, the diffusion dynamics does not affect the separation of IFCs and only influences quantitative determination of the molecular brightness and the absolute concentrations (see Fig. S1b,d,e). Therefore, a relatively long $T_{\mathrm{bin}}$ may be used when intramolecular dynamics is negligible. Nevertheless, if necessary, the effect of finite bin-size due to the molecular diffusion on the cumulants of fluorescence photons can be quantitatively estimated using previously reported methods.[5, 6]

Another issue that potentially deteriorates the performance of IFCA is detector nonideality, i.e., detection dead time and afterpulsing, which artificially modulates $I_x(T)$ in the nano- to microsecond time scale.[7, 8] As a workaround for this problem, we use only photon triplets and photon pairs in which all photon time intervals are longer than a threshold value $T_{\mathrm{g}}$ in the computation of $\boldsymbol{\mathcal{T}}$ and $\mathbf{M}$, respectively. This selection process results in a decrease in the number of photon triplets and photon pairs, which eventually affects the accuracy of calculations in Eqs. (3, 5). To compensate for this effect, the obtained tensors, $\boldsymbol{\mathcal{T}}^{\mathrm{g}}$ and $\mathbf{M}^{\mathrm{g}}$, are rescaled as

$$\mathcal{T}_{ijk} = \mathcal{T}_{ijk}^{\mathrm{g}} \times \frac{T_{\mathrm{bin}}^3}{(T_{\mathrm{bin}} - 2T_{\mathrm{g}})(T_{\mathrm{bin}} - 2T_{\mathrm{g}} - T_{\mathrm{c}})(T_{\mathrm{bin}} - 2T_{\mathrm{g}} - 2T_{\mathrm{c}})}, \tag{S1}$$

$$M_{ij} = M_{ij}^{\mathrm{g}} \times \frac{T_{\mathrm{bin}}^2}{(T_{\mathrm{bin}} - T_{\mathrm{g}})(T_{\mathrm{bin}} - T_{\mathrm{g}} - T_{\mathrm{c}})}. \tag{S2}$$

Here, $T_{\mathrm{c}}$ indicates the macrotime clock period and is included for reflecting the fact that two photons cannot possess the same macrotimes.

##### 2. Derivation of fluorescence patterns and concentrations

The scaled second-order cumulant tensor $\mathbf{M}'$ (Eq. (7) in the main text) is represented as a sum over species contained in the sample:

$$\mathbf{M}' = \sum_{s=1}^{n} \mathbf{M}'^{s}, \tag{S3}$$

$$M_{ij}'^{s} = \frac{d_i^s d_j^s N_s}{\sqrt{v_i v_j}}. \tag{S4}$$

By introducing two matrices, $\mathbf{D}': D_{is}' \equiv d_i^s/\sqrt{v_i}$ and $\mathbf{N}: N_{ss'} \equiv N_s \delta_{ss'}$, we can rewrite the above expression as

$$\mathbf{M}' = \mathbf{D}'\mathbf{N}\mathbf{D}'^{\mathrm{T}}. \tag{S5}$$

The eigenvalue decomposition of $\mathbf{M}'$ is

$$\mathbf{M}' = \mathbf{B}\mathbf{W}\mathbf{B}^{\mathrm{T}}. \tag{S6}$$

This can be approximated by using a $x_0 \times n$ matrix $\mathbf{B}': B'_{ip} = B_{ip}$ $(1 \le i \le x_0$ and $1 \le p \le n)$ and a $n \times n$ matrix $\mathbf{W}': W'_{pq} = W_{pq}$ $(1 \le p, q \le n)$:

$$\mathbf{M}' \approx \mathbf{B}'\mathbf{W}'\mathbf{B}'^{\mathrm{T}}. \tag{S7}$$

For Eqs. (S5) and (S7) to be equal, it is necessary that

$$\mathbf{D}'\mathbf{N}^{\frac{1}{2}} = \mathbf{B}'\mathbf{W}'^{\frac{1}{2}}\mathbf{U}^{\mathrm{T}}, \tag{S8}$$

where $\mathbf{U}$ is an arbitrary orthogonal matrix, i.e., $\mathbf{U}^{\mathrm{T}}\mathbf{U} = \mathbf{I}$ ($\mathbf{I}$ is the identity matrix). From this relation, it follows

$$\mathbf{D}' = \mathbf{B}'\mathbf{W}'^{\frac{1}{2}}\mathbf{U}^{\mathrm{T}}\mathbf{N}^{-\frac{1}{2}} \tag{S9}$$

or

$$\frac{d_i^s}{\sqrt{v_i}} = \sum_{p=1}^{n} B_{ip}\sqrt{W_p}U_{sp}\frac{1}{\sqrt{N_s}}. \tag{S10}$$

The scaled third-order cumulant tensor (Eq. (6) in the main text) is represented as the sum over species as

$$\mathcal{T}'_{ijk} = \frac{1}{\sqrt{v_i v_j v_k}}\sum_{s=1}^{n} d_i^s d_j^s d_k^s N_s\,. \tag{S11}$$

By substituting Eq. (S10) into this equation, we obtain

$$\begin{aligned}\mathcal{T}'_{ijk} &= \sum_{s=1}^{n}\left(\sum_{p=1}^{n} B_{ip}\sqrt{W_p}U_{sp}\frac{1}{\sqrt{N_s}}\right)\left(\sum_{q=1}^{n} B_{jq}\sqrt{W_q}U_{sq}\frac{1}{\sqrt{N_s}}\right)\left(\sum_{r=1}^{n} B_{kr}\sqrt{W_r}U_{sr}\frac{1}{\sqrt{N_s}}\right)N_s \\ &= \sum_{s=1}^{n}\left(\sum_{p=1}^{n} B_{ip}\sqrt{W_p}U_{sp}\right)\left(\sum_{q=1}^{n} B_{jq}\sqrt{W_q}U_{sq}\right)\left(\sum_{r=1}^{n} B_{kr}\sqrt{W_r}U_{sr}\right)h_s\,,\end{aligned} \tag{S12}$$

$$h_s = \frac{1}{\sqrt{N_s}}. \tag{S13}$$

By comparing this expression and the experimentally obtained third-order tensor $\boldsymbol{\mathcal{T}}'$, one can determine $\mathbf{U}$ and $h_s$. Using them, the fluorescence pattern of the *s*-th species is obtained as

$$d_i^s = \sqrt{v_i}\sum_{p=1}^{n} B_{ip}\sqrt{W_p}U_{sp}h_s\,, \tag{S14}$$

and its concentration is

$$N_s = \frac{1}{h_s^2}. \tag{S15}$$

##### 3. Jacobi-type algorithm for determination of $\boldsymbol{h_1, \cdots, h_n}$ and U

In IFCA, the third-order cumulant tensor $\boldsymbol{\mathcal{T}}'$ (Eq. (6)) is decomposed into independent components as Eq. (9). In principle, this is accomplished by finding the $h_1, \cdots, h_n$ values and $\mathbf{U}$ in Eq. (9) that best reproduce the experimentally obtained $\boldsymbol{\mathcal{T}}'$. In other words, we try to fit Eq. (6), which represents the experimentally obtained data, with Eq. (9), which corresponds to a prediction by the theory of ICA. In the actual analysis, instead of directly fitting $\boldsymbol{\mathcal{T}}'$, firstly its dimension is reduced by multiplying $\mathbf{B}'$:

$$\mathcal{T}''_{pqr} \equiv \sum_{i=1}^{x_0} \sum_{j=1}^{x_0} \sum_{k=1}^{x_0} B_{ip} B_{jq} B_{kr} \mathcal{T}'_{ijk}, \tag{S16}$$

and then it is compared with its component decomposition:

$$\mathcal{T}''_{pqr} \approx \sum_{s=1}^{n} \sqrt{W_p W_q W_r} U_{sp} U_{sq} U_{sr} h_s. \tag{S17}$$

The optimization process to find the best $h_1, \cdots, h_n$ and $\mathbf{U}$ is carried out as follows.

(i) Choose a random $n \times n$ orthogonal matrix $\mathbf{U}$.

(ii) Set $p = 1, q = 2$.

(iii) Construct a Givens rotation matrix $\mathbf{G}$ using an angle parameter $\theta$, where the $(p,p), (q,p), (p,q), (q,q)$ elements are $\cos\theta, \sin\theta, -\sin\theta, \cos\theta$, respectively, whereas all the other elements are the same as the $n \times n$ identity matrix.

(iv) Replace $\mathbf{U}$ with $\mathbf{GU}$ and search for the best $h_1, \cdots, h_n$ and $\theta$ values that minimize the deviation between Eqs. (S16) and (S17) defined as follows:

$$S = \sum_{p=1}^{n} \sum_{q=1}^{n} \sum_{r=1}^{n} \left( \sum_{i=1}^{x_0} \sum_{j=1}^{x_0} \sum_{k=1}^{x_0} B_{ip} B_{jq} B_{kr} \mathcal{T}'_{ijk} - \sum_{s=1}^{n} \sqrt{W_p W_q W_r} U_{sp} U_{sq} U_{sr} h_s \right)^2. \tag{S18}$$

(v) Update $\mathbf{U}$ with $\mathbf{GU}$ using the obtained best $\theta$ value.

(vi) Repeat (iii)-(v) by changing $p$ from 1 to $n-1$ and $q$ from $p+1$ to $n$.

(vii) Repeat (ii)-(vi) until $S$ reaches a sufficiently small value.

Through this procedure, $h_1, \cdots, h_n$ and $\mathbf{U}$ in Eq. (9) are determined. However, sometimes it may result in a suboptimal solution due to trapping in a local minimum in the optimization process. For avoiding this effect, the whole procedure may be repeated several times by changing the initial choice of a random matrix $\mathbf{U}$ and taking the best result with the smallest $S$ value.

##### 4. Details of IFCA analysis

The photon data of the mixture of fluorescent dyes obtained with the above PIE-TCSPC setup was loaded by using the *bhreader* module from a Python package *phconvert*.[9] In this experiment, the sample was photoexcited by two interleaving pulse trains and the fluorescence signal was detected at two wavelengths, i.e., $c_0 = 2$. In the data analysis, 32 adjacent microtime channels in 4096 raw channels (6.1 ps/ch) were binned together, resulting in $t_0$ = 128 channels (0.195 ns/ch). Consequently, the fluorescence parameter $x$ in IFCA consists of two parts; the first ($x$ = 1-128) and second ($x$ = 129-256) regions correspond to the microtime $t$ of the photons detected at $\lambda_{em}$ = 585 nm and 685 nm, respectively (Fig. 2a). The bin width $T_{bin}$ was set to 5 μs and the data accumulation time was ~1 hr for each data set. The minimum photon interval $T_g$ and the clock period $T_c$ were 101 ns and 50.5 ns, respectively. Note that the afterpulsing effect is practically negligible in the detectors used in this experiment. Three independent data sets were obtained from the same sample and analyzed by IFCA, among which one is presented in Fig. 2a-e,S2 and the other two are shown in Fig. S1f,g. The fluorescence patterns of individual fluorescent dyes (Fig. 2e, blue lines) were obtained by applying IFCA to the reference samples containing only single dyes, where we set $n$ = 2 for each sample and only the predominant component was selected to remove the contributions of background and impurities. All analyses of IFCA were performed by using a custom code written in Python.

The photon data of FRET-labeled single-stranded DNA construct were obtained from a public repository, Zenodo.[10] For IFCA analysis, photon data were loaded by using the *pqreader* module from the *phconvert* package.[9] The microtime resolution of original data, 1280, was reduced by binning 10 adjacent channels to make $t_0$ = 128 (0.16 ns/ch). The first half of the fluorescence pattern ($x$ = 1-128) is from the donor detector and the second half ($x$ = 129-256) is from the acceptor detector. The bin width $T_{bin}$ was set to 20 μs. The minimum photon interval $T_g$ and the clock period $T_c$ were 2 μs and 20 ns, respectively. The number of components was fixed to $n$ = 2. The analysis was repeated for eight data sets recorded with 600 s accumulation time for each. The results of EVD analysis (Eq. 8) and the fitting to third-order cumulant tensor (Eqs. 10 and 11) are shown for the first data set in Fig. 3c,d,S3. From the fluorescence patterns of two components ($d_x^1$ and $d_x^2$), we calculated the apparent FRET efficiency $E$ and the mean donor lifetime $\tau_{donor}$ as follows:

$$E^s = \frac{\sum_{x=129}^{256} d_x^s}{\sum_{x=1}^{256} d_x^s},$$

$$\tau_{\mathrm{donor}}^s = \left[\frac{\sum_{x=1}^{128} x d_x^s}{\sum_{x=1}^{128} d_x^s} - x_{\mathrm{IRF}}\right] \times 0.16 \text{ ns},$$

where $x_{\mathrm{IRF}}$ is the time origin defined as the first moment of the instrument response function.

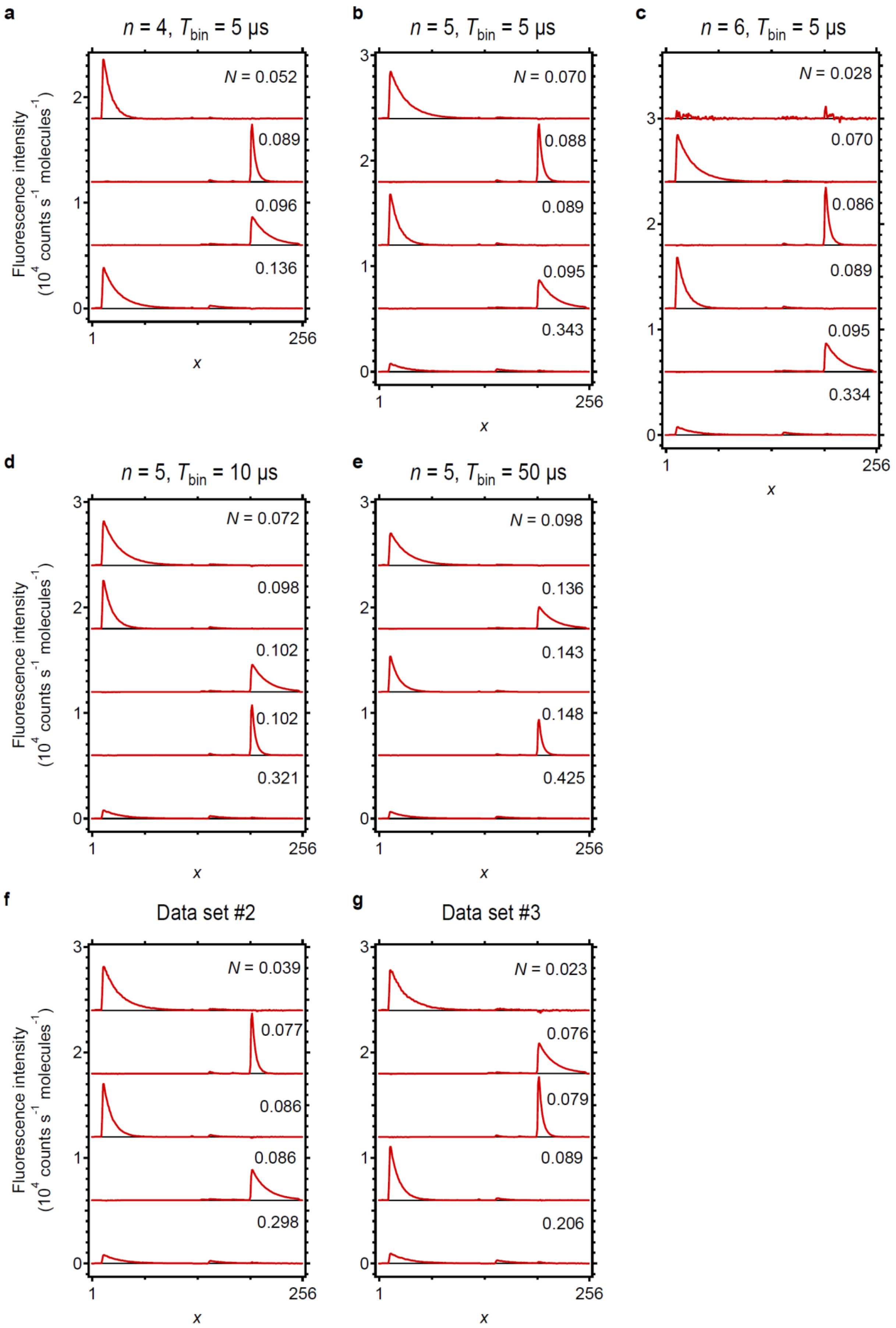
a
n = 4, Tbin = 5 µs
N = 0.052
0.089
0.096
0.136
b
n = 5, Tbin = 5 µs
N = 0.070
0.088
0.089
0.095
0.343
c
n = 6, Tbin = 5 µs
N = 0.028
0.070
0.086
0.089
0.095
0.334
d
n = 5, Tbin = 10 µs
N = 0.072
0.098
0.102
0.102
0.321
e
n = 5, Tbin = 50 µs
N = 0.098
0.136
0.143
0.148
0.425
f
Data set #2
N = 0.039
0.077
0.086
0.086
0.298
g
Data set #3
N = 0.023
0.076
0.079
0.089
0.206
Fluorescence intensity
(10^4 counts s^-1 molecules^-1)
x
1
256

**Fig. S1.** Validation of IFCA on the dye mixture. (a-c) Fluorescence patterns of IFCs obtained with different choices of the number of components, $n$. Binning width was set to $T_{\mathrm{bin}}$ = 5 μs. (d,e) Fluorescence patterns of IFCs obtained with different binning widths ($n$ = 5). (f,g) Fluorescence patterns of IFCs obtained with other data sets ($T_{\mathrm{bin}}$ = 5 μs and $n$ = 5). In all panels, traces are vertically offset for clarity. $N$ values are the average number of molecules in the observation volume estimated by Eq. 13.

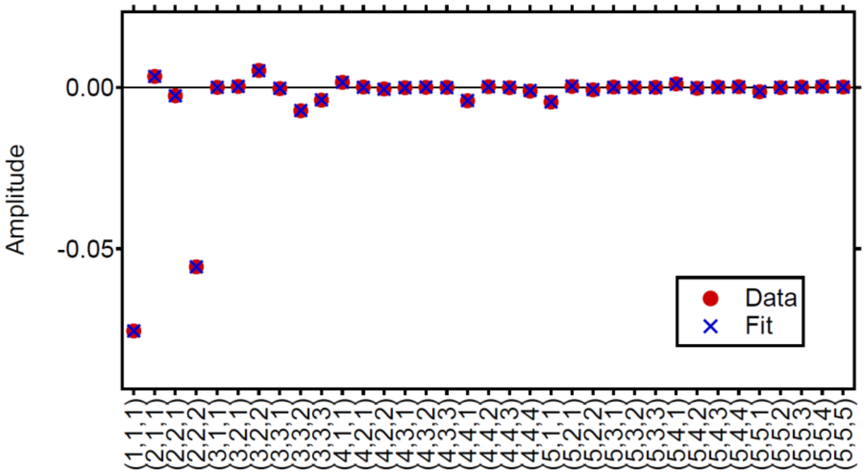


**Fig. S2.** Fitting analysis of the reduced third-order cumulant tensor: Dye mixture. Elements of the tensor are shown individually. Red filled circles: Amplitudes calculated from photon data using Eq. (10). Blue crosses: Fitted values based on Eq. (11).

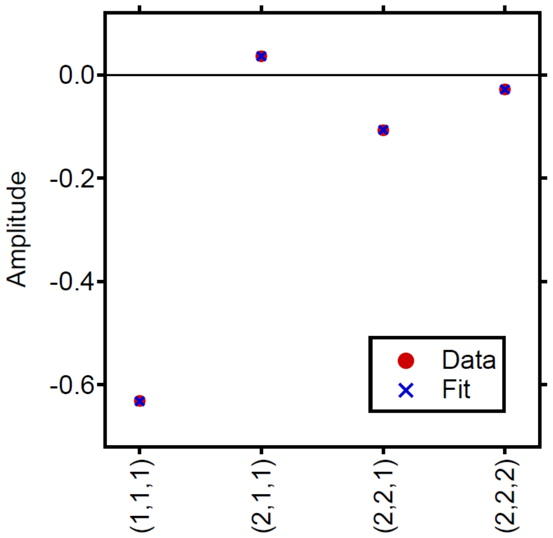


**Fig. S3.** Fitting analysis of the reduced third-order cumulant tensor: FRET-labeled DNA construct. Elements of the tensor are shown individually. Red filled circles: Amplitudes calculated from photon data using Eq. (10). Blue crosses: Fitted values based on Eq. (11).

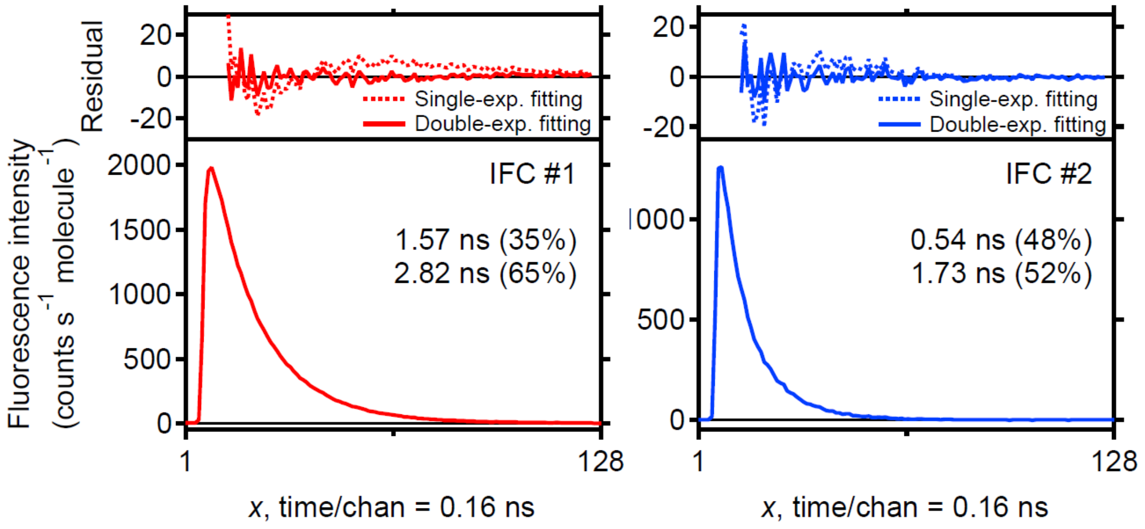


**Fig. S4.** Fitting analysis of the donor decays in two IFCs from the FRET-labeled DNA construct. Decays in the bottom panels are the donor fluorescence ($x$ = 1-128) of two IFCs (left: IFC #1, right: IFC #2). Top panels show fitting residuals derived from single-exponential (dotted line) and double-exponential (solid line) fitting analyses.
